# Speech understanding oppositely affects acoustic and linguistic neural tracking in a speech rate manipulation paradigm

**DOI:** 10.1101/2022.02.04.479105

**Authors:** Eline Verschueren, Marlies Gillis, Lien Decruy, Jonas Vanthornhout, Tom Francart

## Abstract

When listening to continuous speech, the human brain can track features of the presented speech signal. It has been shown that neural tracking of acoustic features is a prerequisite for speech understanding and can predict speech understanding in controlled circumstances. However, the brain also tracks linguistic features of speech, which may be more directly related to speech understanding. We investigated acoustic and linguistic speech processing as a function of varying speech understanding by manipulating the speech rate. In this paradigm, acoustic and linguistic speech processing are affected simultaneously but in opposite directions: When the speech rate increases, more acoustic information per second is present. In contrast, the tracking of linguistic information becomes more challenging when speech is less intelligible at higher speech rates. We measured the EEG of 18 participants (4 male) who listened to speech at various speech rates. As expected and confirmed by the behavioral results, speech understanding decreased with increasing speech rate. Accordingly, linguistic neural tracking decreased with increasing speech rate, but acoustic neural tracking increased. This indicates that neural tracking of linguistic representations can capture the gradual effect of decreasing speech understanding. In addition, increased acoustic neural tracking does not necessarily imply better speech understanding. This suggests that, although more challenging to measure due to the low signal-to-noise ratio, linguistic neural tracking may be a more direct predictor of speech understanding.

**Significance statement:** An increasingly popular method to investigate neural speech processing is to measure neural tracking. Although much research has been done on how the brain tracks acoustic speech features, linguistic speech features have received less attention. In this study, we disentangled acoustic and linguistic characteristics of neural speech tracking via manipulating the speech rate. A proper way of objectively measuring auditory and language processing paves the way towards clinical applications: An objective measure of speech understanding would allow for behavioral-free evaluation of speech understanding, which allows to evaluate hearing loss and adjust hearing aids based on brain responses. This objective measure would benefit populations from whom obtaining behavioral measures may be complex, such as young children or people with cognitive impairments.

## 1 INTRODUCTION

Understanding speech relies on the integration of different acoustic and linguistic properties of the speech signal. The acoustic properties are mainly related to sound perception, while the linguistic properties are linked to the content and understanding of speech. When listening to continuous speech, our brain can track both the acoustic and linguistic properties of the presented speech signal.

Neural tracking of acoustic properties of natural speech has been the subject of many studies. Particular emphasis has been placed on recovering the temporal envelope, i.e., the slow modulations of the speech signal, from the brain responses, so-called neural envelope tracking. The temporal envelope is essential for speech understanding (Shannon et al., 1995), and neural envelope tracking can be linked to speech intelligibility (e.g. Ding and Simon, 2013; Vanthornhout et al., 2018; Lesenfants et al., 2019; Iotzov and Parra, 2019; Verschueren et al., 2020). However, only taking acoustic speech properties into account to investigate neural speech tracking would underestimate the complexity of the human brain, where linguistic properties also play an essential part, as reviewed in detail by Brodbeck and Simon (2020).

In addition to acoustic properties, there is growing interest in retrieving linguistic properties from brain responses to speech. Broderick et al. (2018) used semantic dissimilarity to quantify the meaning carried by words based on their preceding context. They report that the brain responds in a time-locked way to the semantic context of each content word. Additionally, neural tracking is also observed to linguistic properties derived from the probability of a given word or phoneme, i.e., word or phoneme surprisal (Brodbeck et al., 2018; Weissbart et al., 2019; Koskinen et al., 2020). Recently Gillis et al. (2021b) combined several linguistic neural tracking measures and evaluated the potential of each measure as a neural marker of speech intelligibility. After controlling for acoustic properties, phoneme surprisal, cohort entropy, word surprisal, and word frequency were significantly tracked. These results show the potential of linguistic representations as a neural marker of speech intelligibility. In addition, this underlines the importance of controlling for acoustic features when investigating linguistic neural processing, as acoustic and linguistic features are often correlated (Brodbeck and Simon, 2020).

We investigated whether neural speech processing can capture the effect of gradually decreasing speech understanding by manipulating the speech rate. In this study, we focused on acoustic and linguistic speech processing. By changing the speech rate, we manipulate acoustic and linguistic speech processing simultaneously but in opposite directions: When increasing the speech rate, more phonemes, words, and sentences, and thus more acoustic information per second is present. In contrast, linguistic tracking decreases because it becomes more challenging to identify the individual phonemes or words at high speech rates, causing decreased speech understanding. We hypothesize that neural tracking of acoustic features will increase with increasing speech rate because more acoustic information will be present. However, linguistic speech tracking will decrease with increasing speech rate because of decreasing speech understanding. The effect of speech rate on neural responses to speech has already been investigated. However, all these studies only investigated brain responses to the acoustic properties of the speech signal (Ahissar et al., 2001; Nourski et al., 2009; Hertrich et al., 2012; Müller et al., 2019; Casas et al., 2021). It is challenging to extrapolate these findings to what would happen at a linguistic level as acoustic aspects of the speech dominate the neural responses to speech. No study, to our knowledge, reported on how speech rate affects linguistic speech processing and the potential interaction between both. In addition, no consensus has been reached on the effect of speech rate on acoustic neural tracking. For example, Nourski et al. (2009) reported that phase-locked responses decrease with increasing speech rate, similar to Ahissar et al. (2001) and Hertrich et al. (2012). However, in the same data, Nourski et al. (2009) also reported that time-locked responses to the envelope (70-250 Hz) could still be found at very high speech rates where speech is no longer understood.

We investigated how linguistic and acoustic speech tracking are affected when speech understanding gradually decreases. Analyzing neural speech tracking to different characteristics of the presented speech allows us to identify neural patterns associated with speech understanding.

## 2 MATERIAL AND METHODS

### 2.1 Participants

Eighteen participants aged between 19 and 24 years (4 men and 14 women) took part in the experiment after having provided informed consent. Participants had Dutch as their mother tongue and were all normal-hearing, confirmed with pure tone audiometry (thresholds *≤* 25 dB HL at all octave frequencies from 125 Hz to 8 kHz). The study was approved by the Medical Ethics Committee UZ Leuven / Research (KU Leuven) with reference S57102.

### 2.2 Speech material

The story presented during the EEG measurement was ‘A casual vacancy’ by J.K. Rowling, narrated in Dutch by Nelleke Noordervliet. The story was manually cut into 12 blocks of varying length selected from the following list: 4 min, 5 min, 8.5 min, 12.5 min, 18 min, and 23 min. After cutting the story, the story was time-compressed with the Pitch Synchronous Overlap and Add algorithm (PSOLA) from PRAAT (Boursma and Weenink, 2018) to manipulate the speech rate. This algorithm compresses the speech by dividing the speech into small overlapping segments and then repeating or deleting segments to decrease or increase the speech rate, thereby preserving the pitch. However, this manipulation is not exactly comparable with fast natural speech where vowels and silences become shorter, while some consonants (like, e.g, ‘t’) cannot be shortened. Using this algorithm all phonemes are merging similarly and the formant transitions are probably not completely realistic anymore. Six different compression ratio’s (CR) were used: 1.4, 1.0, 0.6, 0.4, 0.28, 0.22 with corresponding speech rates varying from ≈ 2.6 syllables/second (CR=1.4) to ≈ 16.2 syllables/second (CR=0.22). An example audio fragment for every speech rate is added as supplementary material. The fastest CR (CR=0.22) was applied to the longest part (23 min), the one but fastest CR (CR=0.28) to the one but longest part (18 min), and so on. This way, all story parts were compressed or expanded to ≈ 5 minutes. These blocks had slightly different lengths because word and sentence boundaries were taken into account while cutting the story, which is important for the linguistic analysis. Every speech rate was presented twice to obtain 10 minutes of speech at the same rate. The story was presented in chronological order and no part of the story was presented twice to the same participant. Although the linguistic measures only go back up to 4 words, and higher top-down processing has limited impact on the linguistic measures we used, we presented all speech rates in random order per participant. This way the effect of the level of understanding of previous parts, which could influence linguistic processing, was minimized. For each stimulus block, we determined the number of syllables using the forced aligner of the speech alignment component of the reading tutor (Duchateau et al., 2009) and CELEX database (Baayen et al., 1996). The number of syllables uttered for each speech block was then divided by the duration of the speech block in seconds to obtain the speech rate.

After each part of the story, content questions were asked to maximize the participants’ attention and motivation. In addition, speech intelligibility was measured after each block by asking the participants to rate their speech understanding on a scale from 0 to 100% following the question ‘Which percentage of the story did you understand?’. A short summary of the story was shown in the beginning of the experiment to enhance intelligibility as some participants started with more difficult speech rates.

### 2.3 Experimental setup

#### 2.3.1 EEG recording

EEG was recorded with a 64-channel BioSemi ActiveTwo EEG recording system at a sample rate of 8192 Hz. Participants sat in a comfortable chair and were asked to move as little as possible during the EEG recordings. All stimuli were presented bilaterally using APEX 4 (Francart et al., 2008), an RME Multiface II sound card (Haimhausen, Germany), and Etymotic ER-3A insert phones (Illinois, USA). The setup was calibrated using a 2 cm^3^ coupler of the artificial ear (Brüel & Kjær 4152, Denmark). Recordings were made in a soundproof and electromagnetically shielded room.

### 2.4 Signal processing

#### 2.4.1 EEG processing

We processed the EEG in 5 consecutive steps. Firstly, we drift-corrected the EEG signals by applying a first-order highpass Butterworth filter with a cutoff frequency of 0.5 Hz in the forward and backward direction. Then, we reduced the sampling frequency of the EEG from 8192 Hz to 256 Hz to reduce computation time. Artifacts related to eyeblinks were removed with a multichannel Wiener filter (Somers et al., 2018). Subsequently, we referenced the EEG signals to the common average signal. The major advantage of common average referencing is that topography patterns are not biased relative to the chosen channel. No particular accentuation of certain channels allows us to investigate the contribution of neural processes related to language processing evoked by spatially different neural sources. Lastly, we removed the power line frequency of 50 Hz by using a second-order IRR notch filter at this frequency with a quality factor of 35 to determine the filter’s bandwidth at the -3dB.

#### 2.4.2 Stimuli Representations

This study aims to investigate acoustic and linguistic neural tracking at different speech rates. To examine acoustic tracking, we estimated neural tracking based solely on acoustic representations of the stimulus, namely the spectrogram and acoustic edges. To investigate linguistic neural tracking, we created two models: a model to control speech acoustics, which consisted of acoustic and lexical segmentation representations, and a model that included linguistic representations on top of these acoustic and lexical segmentation representations. All speech representations used in the analysis are visualized in Figure 1.

**Figure 1.**
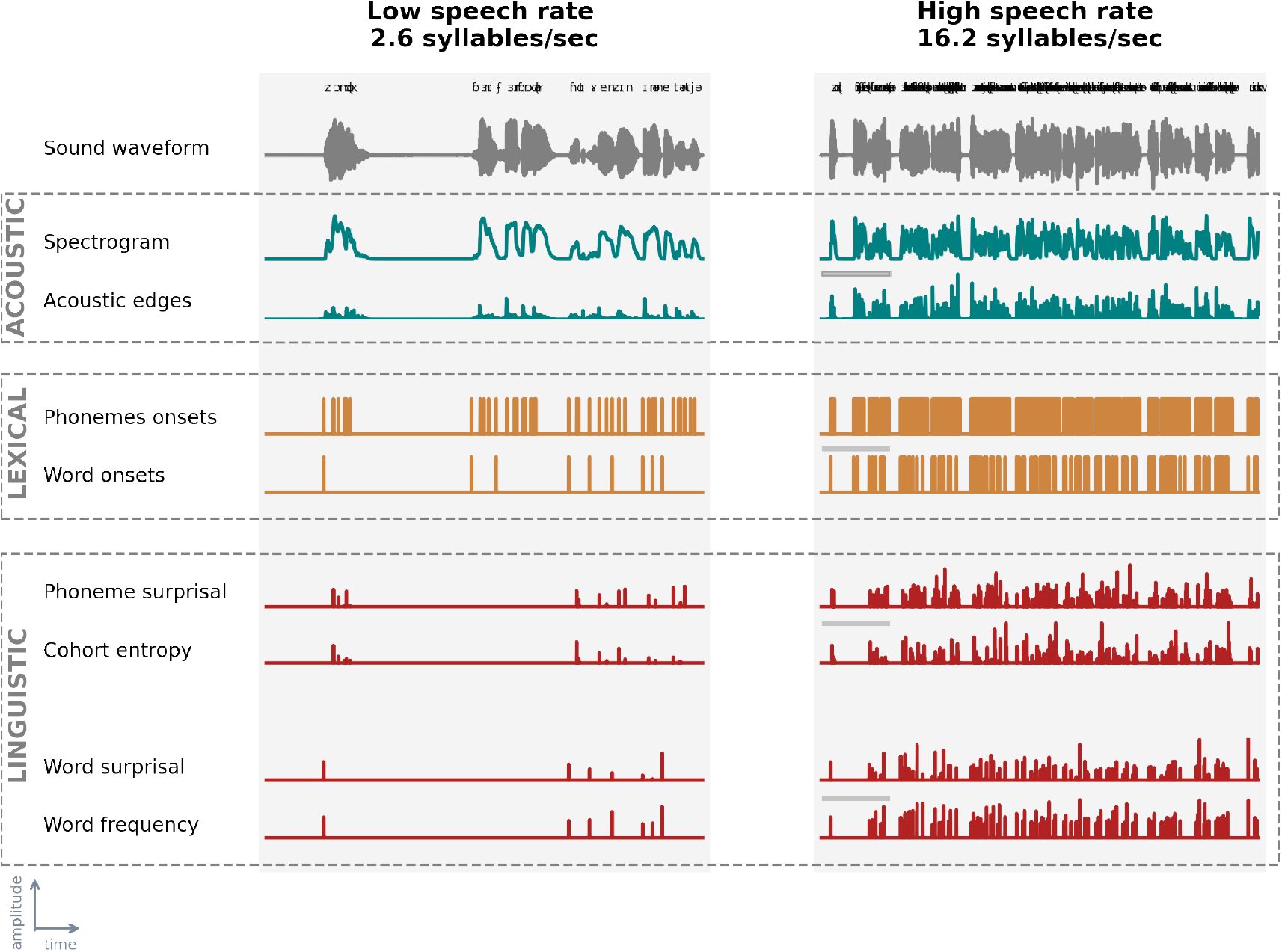
Speech representations at acoustic, lexical and linguistic level. We visualized the speech representations used in this study for all three levels: acoustic (averaged across frequency bands; top box; blue), lexical (middle box; orange), and linguistic (bottom box; red) for the lowest speech rate (SR = 2.6 syllables/sec, left) and highest speech rate (SR = 16.2 syllables/sec; right) for the first 10 seconds of the speech material. This is the first sentence of the audio book. Note: Barry Fairbrother (second and third word) is a proper name and therefore not included in the corpus used for the linguistic features.

The spectrogram representations were calculated based on the low-pass filtered speech stimulus (zero-phase low-pass FIR filter with a hamming window of 159 samples). We low-pass filtered the stimulus at a cut-off frequency of 4 kHz as the insert earphones also low-pass filter at this frequency. Subsequently, we calculated the spectrogram representation from this filtered stimulus using the Gammatone Filterbank Toolkit (Heeris, 2014, center frequencies between 70 and 4000 Hz with 256 filter channels and an integration window of 0.01 second). By using a Gammatone Filterbank, the estimated filter outputs are closer to the human auditory response: the higher the frequency, the wider the filter (Slaney, 1998). We combined the filter outputs by averaging them into eight frequency bands with center frequencies of 124 Hz, 262 Hz, 455 Hz, 723 Hz, 1098 Hz, 1618 Hz, 2343 Hz, and 3352 Hz. To calculate the acoustic edges representations, we took the derivative of the spectrogram’s response in each frequency band and mapped all its negative values to 0. Lastly, we reduced the sampling frequency of these representations to the same sampling frequency as the EEG, namely 256 Hz.

To determine linguistic neural tracking, we carefully controlled for neural responses related to acoustic and lexical characteristics of the speech. As pointed out by Brodbeck and Simon (2020); Gillis et al. (2021b), it is important to control for these characteristics when investigating linguistic neural tracking, as otherwise spurious linguistic neural tracking can be observed due to the high correlation between linguistic and acoustic representations. To evaluate linguistic neural tracking, we determined the added value of linguistic representations by subtracting the performance of the model containing acoustic and lexical segmentation characteristics of the speech from the performance of the model that included the same representations together with the linguistic representations. We used four linguistic representations: phoneme surprisal, cohort entropy, word surprisal, and word frequency, which according to Gillis et al. (2021b), have an added value over and beyond acoustic representations.

All linguistic representations are one-dimensional arrays with impulses at the onsets of phonemes or words. The amplitude of an impulse represents the amount of linguistic information conveyed by the phoneme or word. To obtain the timing of the phonemes and words, we used the forced aligner of the speech alignment component of the reading tutor (Duchateau et al., 2009). Similar to linguistic representations, lexical segmentation representations are one-dimensional arrays. However, the impulses’ amplitudes are one and thus independent of the amount of linguistic information.

Phoneme surprisal and cohort entropy are two metrics that summarise upcoming speech content’s information value and expected information value. Phoneme surprisal is calculated as the negative logarithm of the inverse conditional probability of the phoneme given the preceding phonemes of the word. It is a measure of phoneme prediction error as it represents how surprising a phoneme is given the previously uttered phonemes (Brodbeck et al., 2018; Gwilliams and Davis, 2022). Therefore, phoneme surprisal reflects to what extent the upcoming phoneme can be predicted, which is intrinsically related to the information contributed by the phoneme. The more likely a specific phoneme is, the lower its surprisal value, and the less information is gained (and vice versa). Cohort entropy is calculated as the Shannon entropy of the active cohort of words, i.e., the cohort of words congruent with the already uttered phonemes. Cohort entropy is a measure of uncertainty: higher cohort entropy implies that the active cohort consists of a larger number of words with a similar probability of being uttered (Brodbeck et al., 2018; Gwilliams and Davis, 2022). Both measures have a similar trend: phoneme surprisal and cohort entropy tend to decrease when more phonemes of the word are pronounced because the prediction of the upcoming word can be made with a higher precision. However, if the upcoming word cannot be predicted with high precision, these metrics at phoneme level can increase.

To calculate both representations, we used a custom pronunciation dictionary that maps a word to its phoneme representation. This dictionary was created by manual and grapheme-to-phoneme conversion and contained the segmentation of 9157 words. The word probabilities were derived from the SUBTLEX-NL database (Keuleers et al., 2010). The linguistic information of the initial phoneme was not modeled in these representations. More details regarding phoneme surprisal and cohort entropy, as well as the mathematical determinations, can be found in Brodbeck et al. (2018).

The linguistic information conveyed by a word is described by word surprisal and word frequency. Similarly as phoneme surprisal, word surprisal reflects to what extent the upcoming word can be predicted based on the information contributed by that word. We used a 5-gram model to determine the negative logarithm of the conditional probability of the word given the preceding words. Therefore, a word’s surprisal is estimated given its four preceding words. Word frequency was derived from the same 5-gram model but without including previous words, describing the word’s unigram probability. This measure reflects the probability of the word, independent of its context.

Notice that these linguistic representations rely on the statistical probabilities in language. Gwilliams and Davis (2022) reviewed in detail how listeners rely on this type of internal language model that allows them to predict the upcoming word or phoneme.

#### 2.4.3 Determination of Neural Tracking

To determine neural tracking, we used a forward modeling approach, estimating how the brain responds to specific speech characteristics. The temporal response function (TRF) describes the relationship between the presented stimulus and measured EEG. It also allows us to predict the EEG responses associated with the speech stimulus. By correlating the predicted EEG responses with the measured EEG responses, we obtain a prediction accuracy per EEG channel. This prediction accuracy is a measure of neural tracking.

We used the boosting algorithm (David et al., 2007) implemented by the Eelbrain Toolbox (Brodbeck, 2020) to estimate the TRF and obtain the prediction accuracy. The boosting algorithm estimates the TRF in a sparse and iterative way. It requires a training partition to estimate the TRF and a validation partition to evaluate the TRF. The boosting algorithm aims to find the optimal solution by evaluating the effect of incremental changes in the TRF. The incremental change which yields the highest prediction accuracy on the validation set, is kept. Incremental changes are made until no further improvement can be found on the validation set.

We used an integration window of -100 to 600 ms, i.e., the neural response is estimated ranging from 100 ms before activation of the stimulus characteristic to 600 ms after its activation. We use a broad integration window to ensure that the model captures the brain responses to the linguistic representations, which occur at longer latencies. As each speech rate condition was presented twice, we estimated the TRF on the concatenation of these two blocks per speech rate, i.e., ten minutes of data. Before the TRF estimation, the data is normalized by dividing by the Euclidean norm per channel. We applied this normalization for the stimulus and EEG data individually. Then the boosting algorithm estimates the associated response behavior using a fixed step size of 0.005. We derived the TRF and prediction accuracy per channel using a cross-validation scheme: the TRF was estimated and validated on partitions unseen during testing of the TRF to obtain the prediction accuracy. More specifically, we used 10-fold cross-validation, implying the data was split into ten equally long folds, of which eight folds are used for estimating the TRF, one fold for validation, and one fold for testing. The obtained TRFs and prediction accuracies are then averaged across the different folds. Note that the validation fold is required to determine the stopping criterion: we used an early stopping based on the *𝓁*_2_-norm, i.e., estimation of the TRF is stopped when the Euclidian distance between the actual and predicted EEG data on the validation partition stops decreasing. The resulting TRFs are sparse. Therefore, to account for the inter-subject variability and obtain a meaningful average TRF response across subjects, we smoothed the TRFs across time by convolving the estimated response with a hamming window of 50 ms in the time dimension.

To determine the acoustic tracking of the speech, we purely used acoustic representations. Therefore, we determined the prediction accuracy and TRFs based on the spectrogram and acoustic edges. Regarding the linguistic tracking of speech, we investigated the added value of these linguistic representations. To determine the added value, we subtracted the prediction accuracies of two different models. Firstly, we estimated a baseline model consisting of acoustic and lexical segmentation representations. Secondly, we estimated a combined model which included linguistic representations on top of the acoustic and lexical segmentation representations. By subtracting the prediction accuracy obtained with the baseline model from the prediction accuracy of the combined model, we can examine the added value of the linguistic representations after controlling for the acoustic and lexical segmentation representations. Please note that we did not include lexical features in the acoustic or linguistic model because the neural response to a phoneme or word onset does not reflect purely acoustic nor linguistic processing (Sanders and Neville, 2003).

We used two predetermined channel selections to investigate the effect of acoustic and linguistic tracking. The neural responses to acoustics are significantly different from those to linguistic content and therefore require a different channel selection. We used a frontal channel selection for acoustic neural tracking based on Lesenfants et al. (2019) and a central channel selection for linguistic neural tracking as reported by Gillis et al. (2021b).

These channel selections were used to visualize the TRFs and to determine associated peak latency and amplitudes. To determine the peak characteristics, we set a preset time window based on the TRF averaged across subjects (see Table 1). Within this time window, we normalized the TRF per channel by dividing the TRF by its *𝓁*_2_-norm over time to decrease across subject variability and averaged the TRF across the channel selection. Depending on a positive or negative peak, we determined the maximal or minimal amplitude and its corresponding latency to obtain the peak amplitude and latency. If the peak latency was the same as the beginning of the window, indicating the end of the previous peak, we discarded the peak from the analysis (see Table 2).

**Table 1.** Time windows selected per speech representation to determine the peak characteristics

| Speech representation | Time window(s) | Channel selection |
| --- | --- | --- |
| Spectrogram<br>Acoustic edges | 0 to 90 ms,<br>110 to 200 ms | frontocentral |
| Phoneme surprisal<br>Cohort entropy | 200 to 300 ms | central |
| Word surprisal<br>Word frequency | 350 to 450 ms | central |

**Table 2.** Number of peaks detected per speech representation per speech rate with n_max_ = 18 (= amount of participants).

|  | 2.6 syll/sec | 3.6 syll/sec | 6.2 syll/sec | 9.0 syll/sec | 12.9 syll/sec | 16.2 syll/sec |
| --- | --- | --- | --- | --- | --- | --- |
| Spectrogram - peak 1 | n = 15 | n = 17 | n = 18 | n = 18 | n = 18 | n = 18 |
| Spectrogram - peak 2 | n = 14 | n = 16 | n = 18 | n = 15 | n = 13 | n = 11 |
| Acoustic edges - peak 1 | n = 17 | n = 17 | n = 18 | n = 18 | n = 18 | n = 18 |
| Acoustic edges - peak 2 | n = 18 | n = 17 | n = 16 | n = 10 | n = 8 | n = 8 |
| Phoneme surprisal | n = 17 | n = 18 | n = 17 | n = 16 | n = 15 | n = 18 |
| Cohort entropy | n = 17 | n = 16 | n = 16 | n = 15 | n = 18 | n = 17 |
| Word surprisal | n = 15 | n = 16 | n = 15 | n = 18 | n = 16 | n = 17 |
| Word frequency | n = 16 | n = 15 | n = 13 | n = 14 | n = 15 | n = 17 |

### 2.5 Statistics

Statistical analysis was performed using MATLAB (version R2018a) and R (version 3.4.4) software. The significance level was set at *α*=0.05 unless otherwise stated.

To evaluate the subjectively rated speech understanding results we calculated the correlation between speech rate and rated speech understanding using a Spearman rank correlation. In addition, we fitted a sigmoid function on the data to address the relation between rated speech understanding and speech rate using the minpack.lm package (Elzhov et al., 2016) in R. For further statistical analysis, we selected speech rate (and not subjectively rated speech understanding) as a main predictor. We opted for this because (1) it is difficult to obtain reliable speech understanding scores from a continuous story. In clinical practice participants are asked to repeat sentences or words to calculate a percentage correct. Asking someone to repeat an entire discourse and scoring it is not feasible (Decruy et al., 2018; MacPherson and Akeroyd, 2014). (2) A subjective rating is very idiosyncratic. This results in a high inter-subject variability: some participants will rate their speech understanding more optimistically than others for the same underlying level of understanding.

To determine whether the topographies or the TRFs were significantly different from zero, we performed non-parametric permutation tests (Maris and Oostenveld, 2007). For the analysis of the acoustic TRFs, we limited the window of interest to the time region between 0 and 200 ms. As speech is more difficult to understand, the latency of the neural responses to acoustic representation increases (Mirkovic et al., 2019; Verschueren et al., 2021; Gillis et al., 2021a). These effects are most prominent in a time region of 0 to 200 ms (Verschueren et al., 2020; Mirkovic et al., 2019; Kraus et al., 2020), explaining the rationale to limit the time window of interest. However, no time window of interest was set to determine the significance of the linguistic TRFs. We are not aware of any studies that assess the effect of linguistic tracking when speech comprehension becomes challenging. Therefore we chose not to specify a time window of interest when investigating the neural responses to linguistic representations. As observed in previous literature, linguistic TRFs are associated with negative responses in central areas. Therefore, we applied this test in a one-sided fashion, i.e., we determined where the TRF was significantly negative.

To asses the relationship between speech rate, neural speech tracking and speech understanding, we created a linear mixed effect (LME) model using the LME4 package (Bates et al., 2015) in R with the following general formula:

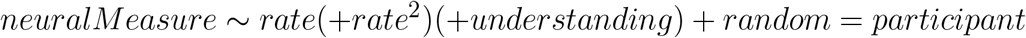

where “neuralMeasure” refers to neural speech tracking, TRF amplitude or TRF latency, depending on the model being investigated and “rate” refers to the speech rate the speech was presented at. Speech rate was also added as a quadratic effect, “rate^2^”, as we do not expect neural speech tracking will decrease or increase linearly indefinitely with increasing speech rate. Lastly, “understanding”, referring to rated speech understanding, was added to the model to investigate whether speech understanding is able to explain additional variance on top of speech rate. An additional random intercept per participant was included in the model to account for the multiple observations per participant. “Rate^2^” and “understanding” are added between brackets to the general formula because these factors were only included if they benefited the model. We controlled this by calculating the Akaike Information Criterion (AIC) for the model with and without “Rate^2^” and “understanding”. The model with the lowest AIC was selected and its residuals plot was analyzed to assess the normality assumption of the LME residuals. Unstandardized regression coefficients (beta) with 95% confidence intervals and p-value of the factors included in the model are reported in the results section.

## 3 RESULTS

### 3.1 Effect of speech rate on speech understanding

Figure 2 shows that when speech rate increases, rated speech understanding decreases (r=-0.91, p*<*0.001, Spearman rank correlation). To model the data, we fitted a sigmoid function between speech understanding and speech rate. The function shows a plateau until 6 syllables/sec (=*±*1.5 times the rate of normal speech). When the speech rate further increases, speech understanding drops. Because speech understanding and speech rate are highly correlated, selecting the one or the other for the analysis should not matter too much. We select speech rate for further analysis in function of neural speech tracking. As mentioned above, speech rate is more reliable than the subjectively rated speech understanding scores since it is objectively derived from the acoustic stimulus.

**Figure 2.**
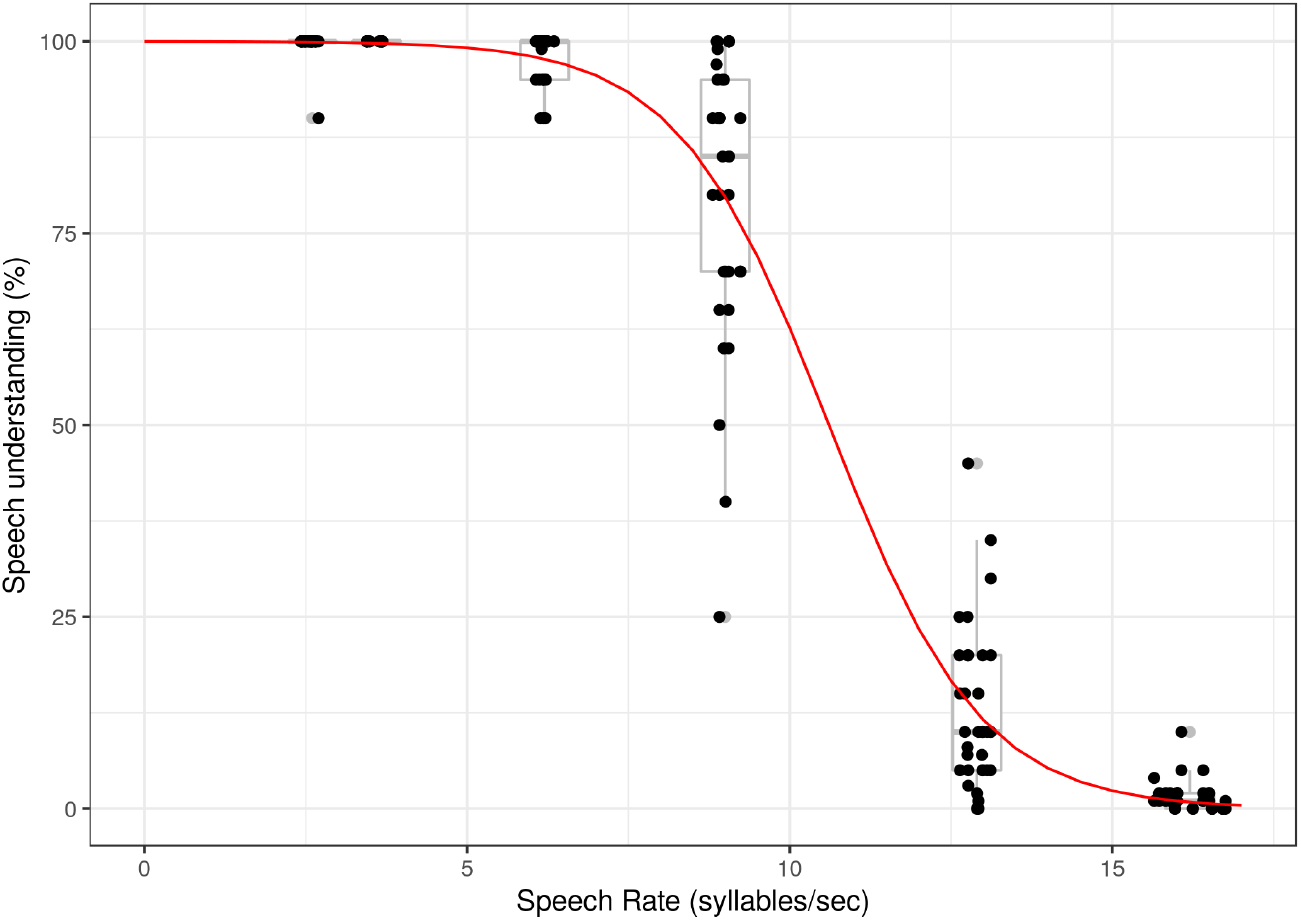
Rated speech understanding in function of speech rate. The dots show speech understanding per participant for every participant specific speech rate. The boxplots show the participants’ results for the averaged speech rates (based on compression ratio). The red line is the sigmoid function fitted on the data over participant.

### 3.2 Effect of speech rate on neural processing of speech

To obtain the results in this section, we created two models: an acoustic model and a linguistic model as explained in detail in section 2.4.3.

#### 3.2.1 Acoustic neural tracking

First, we investigated the acoustic model containing acoustic edges and the spectrogram. Figure 3.A shows how accurately the acoustic model can predict the speech signal for every electrode used. To better quantify this result, we selected the frontocentral channels based on Lesenfants et al. (2019) (channels are highlighted in red in the inset of figure 3.B) and averaged them per subject. This resulted in one neural tracking value per speech rate per subject. Neural tracking of this frontocentral channel selection increased with increasing speech rate (p*<*0.001, b=7.61×10^−3^, CI(95%): *±*1.59×10^−3^, LME, table 3). However, as visualized on the right in figure 3.A this increase is not monotonous, but quadratic (p*<*0.001, b=-3.28×10^−4^, CI(95%): *±*8.41×10^−5^, LME, table 3). Finally, adding speech understanding as an extra predictor to the model does not improve the model (AIC_SR_ = -623, AIC_SR + understanding_ = -605).

**Table 3.**
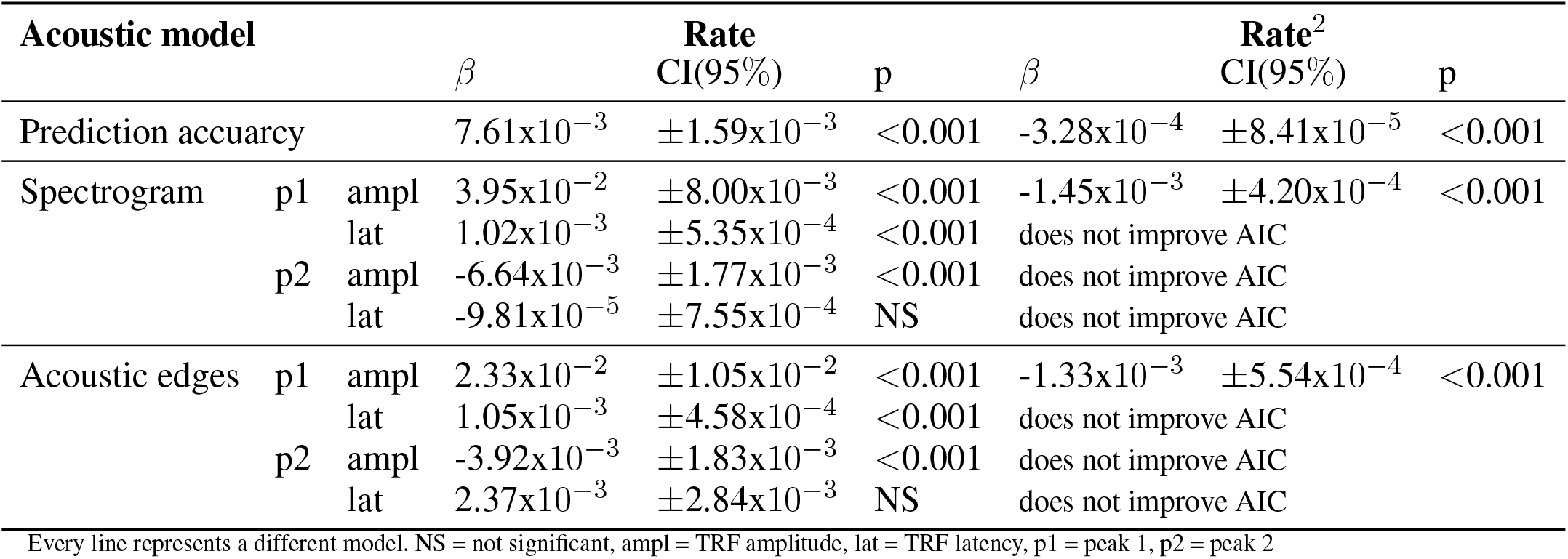
Linear Mixed Effect Model of prediction accuracies and amplitude and latency of the TRF peaks in function of speech rate for the acoustic model

**Figure 3.**
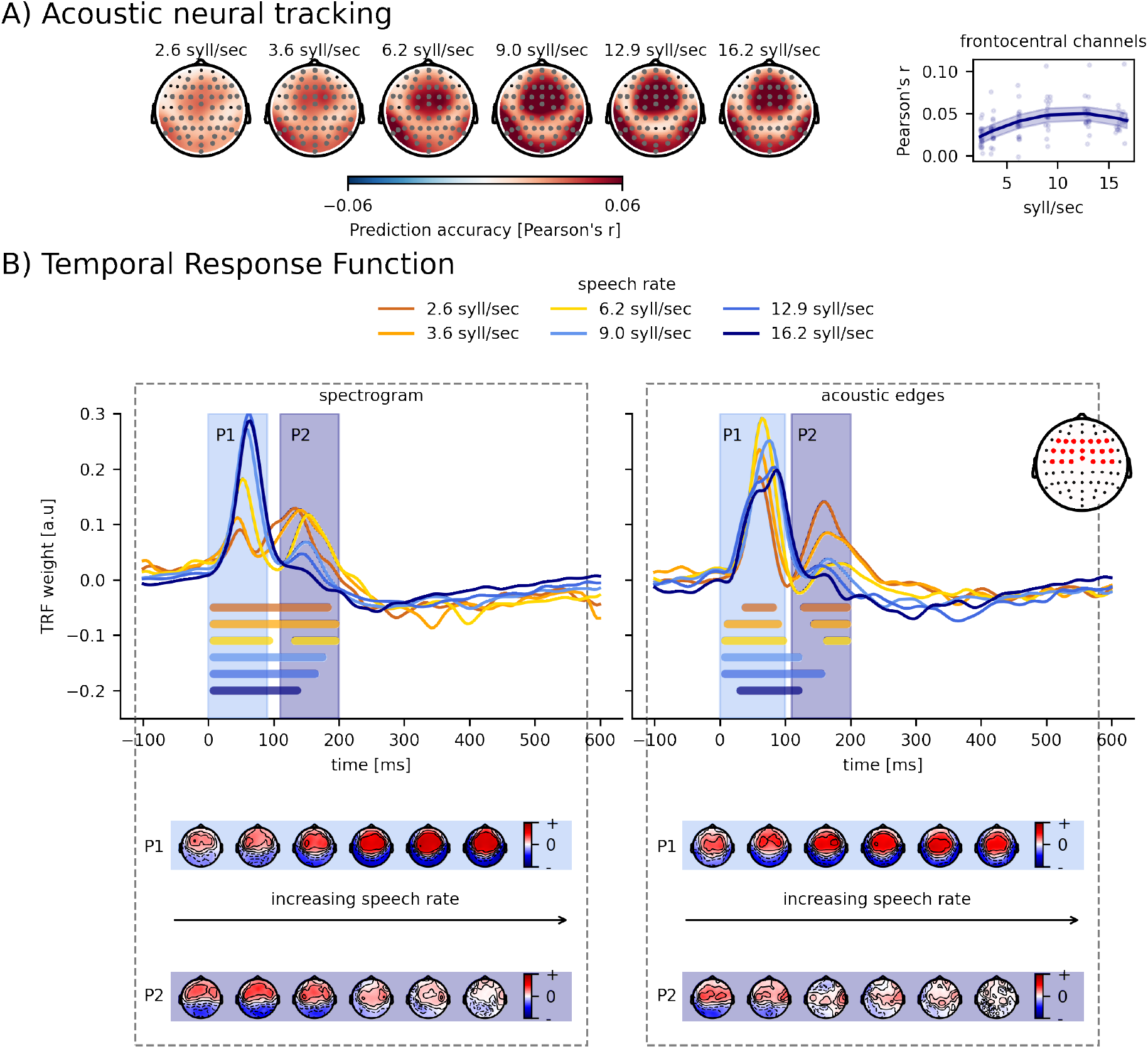
Effect of speech rate on acoustic tracking. Panel A: visualization of the average prediction accuracy across participants for each speech rate. The annotated grey channels indicate the cluster which drives the significant difference from 0. How acoustic tracking, averaged across frontocentral channels, changes according to the speech rate is shown on the right. Panel B: Normalized TRFs of the spectrogram and acoustic edges. The bold horizontal lines indicate where the TRFs are significantly different from 0 (the same color as the TRF of the considered speech rate). The topographies below show the associated peak topographies in the TRF.

To better understand the obtained quadratic tendency of neural tracking as a function of speech rate, we analyzed the acoustic features separately using TRFs. Figure 3.B visualizes the averaged TRF of the frontocentral channels for the spectrogram (left panel) and for the acoustic edges (right panel). For both speech features, two significant positive peaks appear around 70 and 150 ms (horizontal bars show the TRF parts significantly different from zero). The topographies of these peaks are shown underneath the TRFs. More detailed analysis on both peaks is done by calculating the maximum value for every participant per speech rate between 0 and 90 ms (= peak value 1) and between 110 and 200 ms (= peak value 2). We investigated the amplitude and the latency of these peak values as a function of speech rate as shown in Figure 4. With increasing speech rate the amplitude of peak 1 increases, saturates and decreases again for the spectrogram feature (SR: p*<*0.001, b=3.95×10^−2^, CI(95%): *±*8.00×10^−3^; SR^2^: p*<*0.001, b=-1.45×10^−3^, CI(95%): *±*4.20×10^−4^; LME; table 3) and acoustic edges (SR: p*<*0.001, b=2.33×10^−2^, CI(95%): *±*1.05×10^−2^; SR^2^: p*<*0.001, b=-1.33×10^−3^, CI(95%): *±*5.54×10^−4^; LME; table 3). In contrast, the amplitude of peak 2 decreases with increasing speech rate (spectrogram: p*<*0.001, b=-6.64×10^−3^, CI(95%): *±*1.77×10^−3^; acoustic edges: p*<*0.001, b=-3.92×10^−3^, CI(95%): *±*1.83×10^−3^; LME; table 3). Interestingly, the second peak for the acoustic edges even disappears when the speech rate is 9 syllables/sec or higher and speech understanding drops below 80% (Figure 3.B, horizontal bars show the TRF parts significantly different from zero). For the latency analysis of peak 2 for acoustic edges, we thus only include the latency of the peaks in the 3 easiest speech rate conditions, as no peaks (and latencies) can be found anymore at higher speech rates. The latency of peak 1, for both speech features, increases with increasing speech rate (spectrogram: p*<*0.001, b=1.02×10^−3^, CI(95%): *±*5.35×10^−4^; acoustic edges: p*<*0.001, b=1.05×10^−3^, CI(95%): *±*4.58×10^−4^; LME; table 3), while the latency of peak 2 shows no significant relation with speech rate (spectrogram: p=0.80, b=-9.81×10^−5^, CI(95%): *±*7.55×10^−4^; acoustic edges: p=0.11, b=2.37×10^−3^, CI(95%): *±*2.84×10^−3^; LME; table 3).

**Figure 4.**
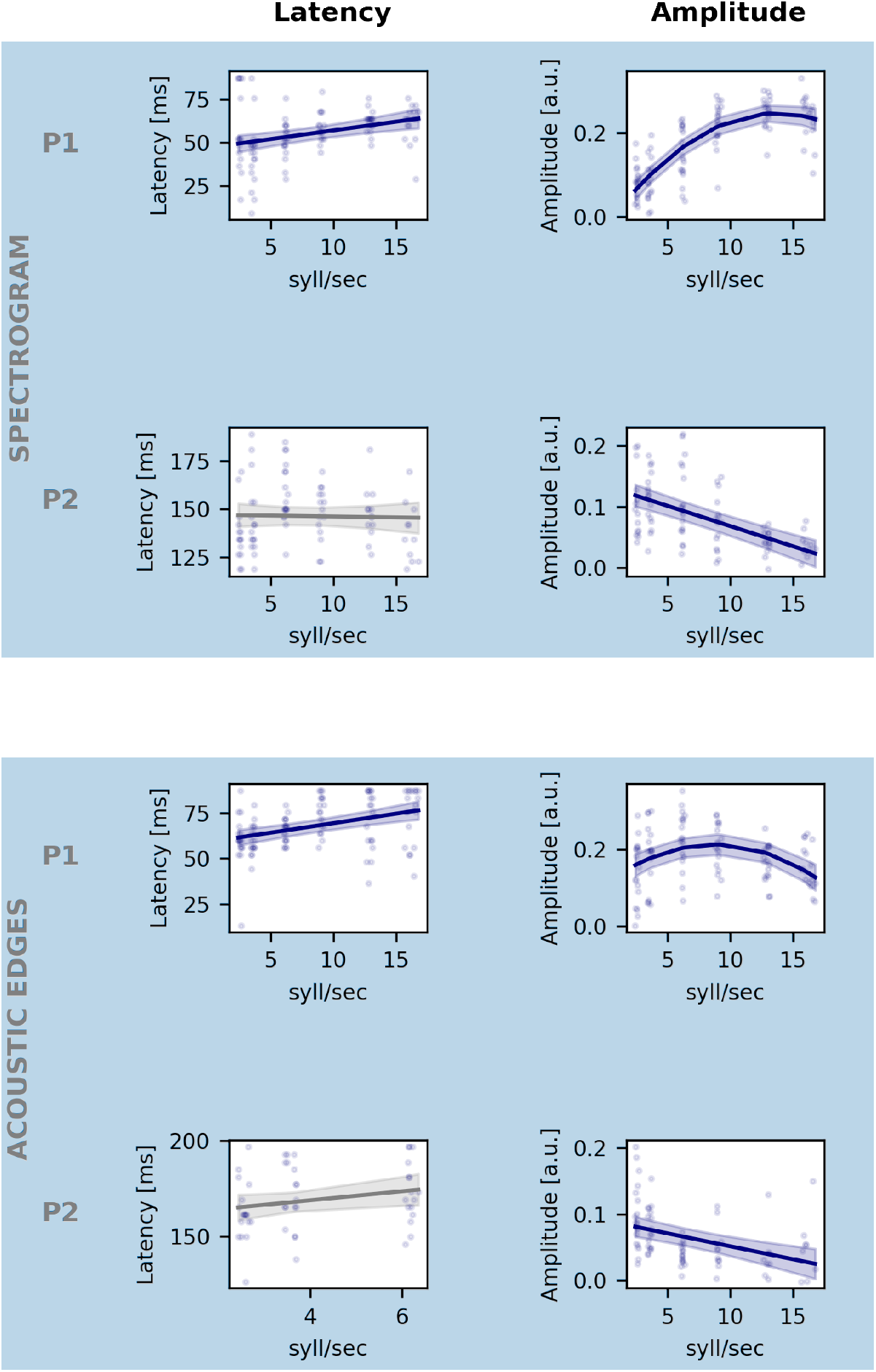
Effect of speech rate on the amplitude and latency of acoustic representation. Each row shows the effect of speech rate on latency (left plot) and the amplitude (right plot) of the neural response to spectrogram (top row) and acoustic edges (bottom row) for respectively the first and second identified peak as indicated on Figure 3. The blue line shows the model’s prediction for each speech rate; the shaded area indicates the confidence interval of the model’s prediction. The non-significant models are shown in grey. Remark that we only include the latency of significant peaks for the latency analysis.

#### 3.2.2 Linguistic neural tracking

Next to acoustic neural tracking, we also investigated the effect of speech rate on linguistic neural tracking (see section 2.4.3 for more details). Figure 5.A (left panel) shows how accurately the linguistic model can predict the speech signal over subjects per speech rate per channel. The channels in the cluster which drives the topography from 0 are annotated with grey markers. The higher the speech rate, the fewer channels have significant neural tracking. To quantify this, we averaged the prediction accuracy over a central channel selection based on Gillis et al. (2021b) (channels are highlighted in red in the inset of figure 5.B), resulting in one neural tracking value per speech rate per subject. As shown in figure 5.A (right), neural tracking significantly drops monotonically with increasing speech rate (p=0.008, b=-4.59×10^−5^, CI(95%):*±*3.31×10^−5^, LME, table 4). Interestingly, this is the opposite trend from the acoustic model in section 3.2.1. Finally, adding speech understanding as a predictor does not improve the linguistic model (AIC_SR_ = -1175, AIC_SR + understanding_ = -1151).

**Table 4.** Linear Mixed Effect Model of prediction accuracy and amplitude and latency of the TRF peaks in function of speech rate for the linguistic model

| Linguistic model | | $\beta$ | rate<br>$CI(95\%)$ | p | $\beta$ | rate<sup>2</sup><br>$CI(95\%)$ | p |
| --- | --- | --- | --- | --- | --- | --- | --- |
| Prediction accuracy | | $-4.59 \times 10^{-5}$ | $\pm 3.31 \times 10^{-5}$ | 0.0081 | does not improve AIC | | |
| Phoneme surprisal | ampl | $6.81 \times 10^{-3}$ | $\pm 2.82 \times 10^{-3}$ | $< 0.001$ | does not improve AIC | | |
| | lat | $5.47 \times 10^{-3}$ | $\pm 4.20 \times 10^{-3}$ | 0.015 | does not improve AIC | | |
| Cohort entropy | ampl | $4.21 \times 10^{-3}$ | $\pm 3.07 \times 10^{-3}$ | 0.008 | does not improve AIC | | |
| | lat | $3.59 \times 10^{-3}$ | $\pm 5.11 \times 10^{-3}$ | NS | does not improve AIC | | |
| Word surprisal | ampl | $6.46 \times 10^{-3}$ | $\pm 3.09 \times 10^{-3}$ | $< 0.001$ | does not improve AIC | | |
| | lat | $-3.33 \times 10^{-3}$ | $\pm 5.38 \times 10^{-3}$ | NS | does not improve AIC | | |
| Word frequency | ampl | $6.44 \times 10^{-3}$ | $\pm 3.00 \times 10^{-3}$ | $< 0.001$ | does not improve AIC | | |
| | lat | $-1.09 \times 10^{-4}$ | $\pm 3.06 \times 10^{-3}$ | NS | does not improve AIC | | |

Every line represents a different model. NS = not significant, ampl = TRF amplitude, lat = TRF latency

**Figure 5.**
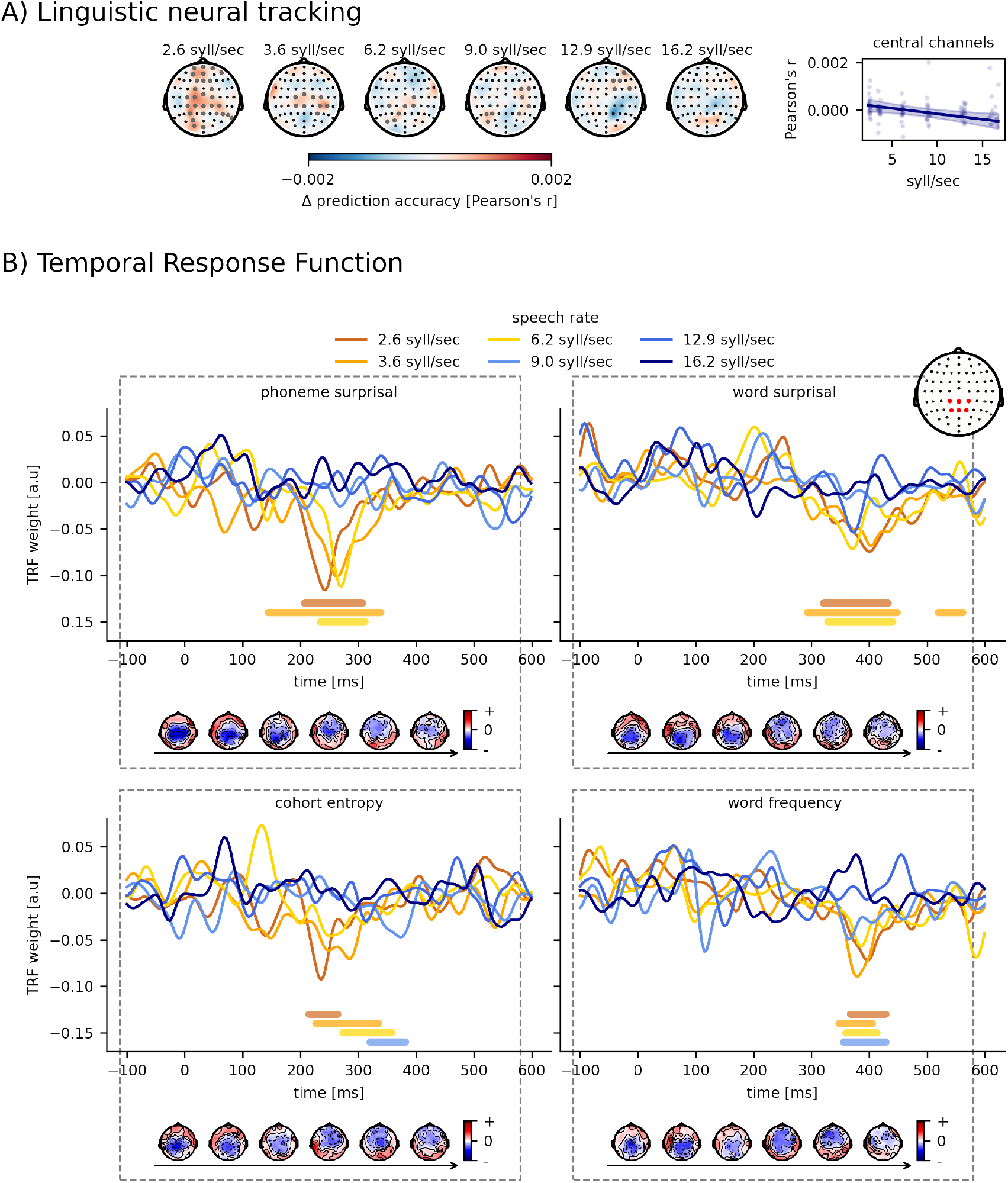
Effect of speech rate on linguistic tracking. Panel A: visualization of the added value of linguistic representations across participants for each speech rate. The annotated grey channels indicate the cluster which drives the significant difference from 0. How linguistic tracking, averaged across central channels, changes according to the speech rate is shown on the right. Panel B: The associated normalized TRFs for the linguistic representations. The bold horizontal lines indicate where the TRFs are significantly different from 0 (the same color as the TRF of the considered speech rate). The topographies below indicate the associated peak topographies to the TRF in the grey shaded area. The horizontal arrow underneath the topographies indicates the increasing speech rate.

To thoroughly investigate the neural responses to the linguistic features, we examined the TRFs of the central channel selection. Figure 5.B visualizes the averaged normalized TRF in the central channel selection for the different linguistic features per speech rate. The grey zone is where, based on Gillis et al. (2021b), we would expect a neural response. Significant responses can be found in the lower speech rates when speech can still be understood for all features. In the higher speech rates, where speech understanding is worse or absent, the linguistic neural response also disappears (Figure 5.B, horizontal bars show the TRF parts significantly different from zero). The topographies of these responses are shown in Figure 5.B underneath the TRFs. Interestingly, the topographies switch from central negativity when speech is understood to frontal negativity when speech understanding is worse or absent. To investigate whether the amplitude or latency of these peaks is related to speech rate, we calculated the minimum value for every participant within the grey zone (= peak value). For all linguistic features the peak amplitude shrinks significantly with increasing speech rate as shown in Figure 6 (Phoneme surprisal: p*<*0.001, b=6.81×10^−3^, CI(95%): *±*2.82×10^−3^; Cohort entropy: p=0.008, b=4.21×10^−3^, CI(95%): *±*3.07×10^−3^; Word surprisal: p*<*0.001, b=6.46×10^−3^, CI(95%): *±*3.09×10^−3^; Word Frequency: p*<*0.001, b=6.44×10^−3^, CI(95%): *±*3.00×10^−3^; LME; table 4). In other words, when speech becomes faster and more difficult to understand, the peak amplitude of the linguistic features decreases until it finally disappears when the speech rate is 9.0 or 12.9 syllables/sec, or higher, and speech understanding is dropping below 90%. Similar to the analysis of the second peak for acoustic features, we only include the latency of significant peaks. Phoneme surprisal shows a significant increase of peak latency with increasing speech rate (p=0.015, b=5.47×10^−3^, CI(95%): *±*4.20×10^−3^; LME; table 4). Cohort entropy, word surprisal and word frequency, on the other hand, reveal no significant effect of speech rate on peak latency (Cohort entropy: p=0.18, b=3.59×10^−3^, CI(95%): *±*5.11×10^−3^; Word surprisal: p=0.23, b=-3.33×10^−3^, CI(95%): *±*5.38×10^−3^; Word frequency: p=0.95, b=-1.09×10^−4^, CI(95%): *±*3.06×10^−3^; LME; table 4).

**Figure 6.**
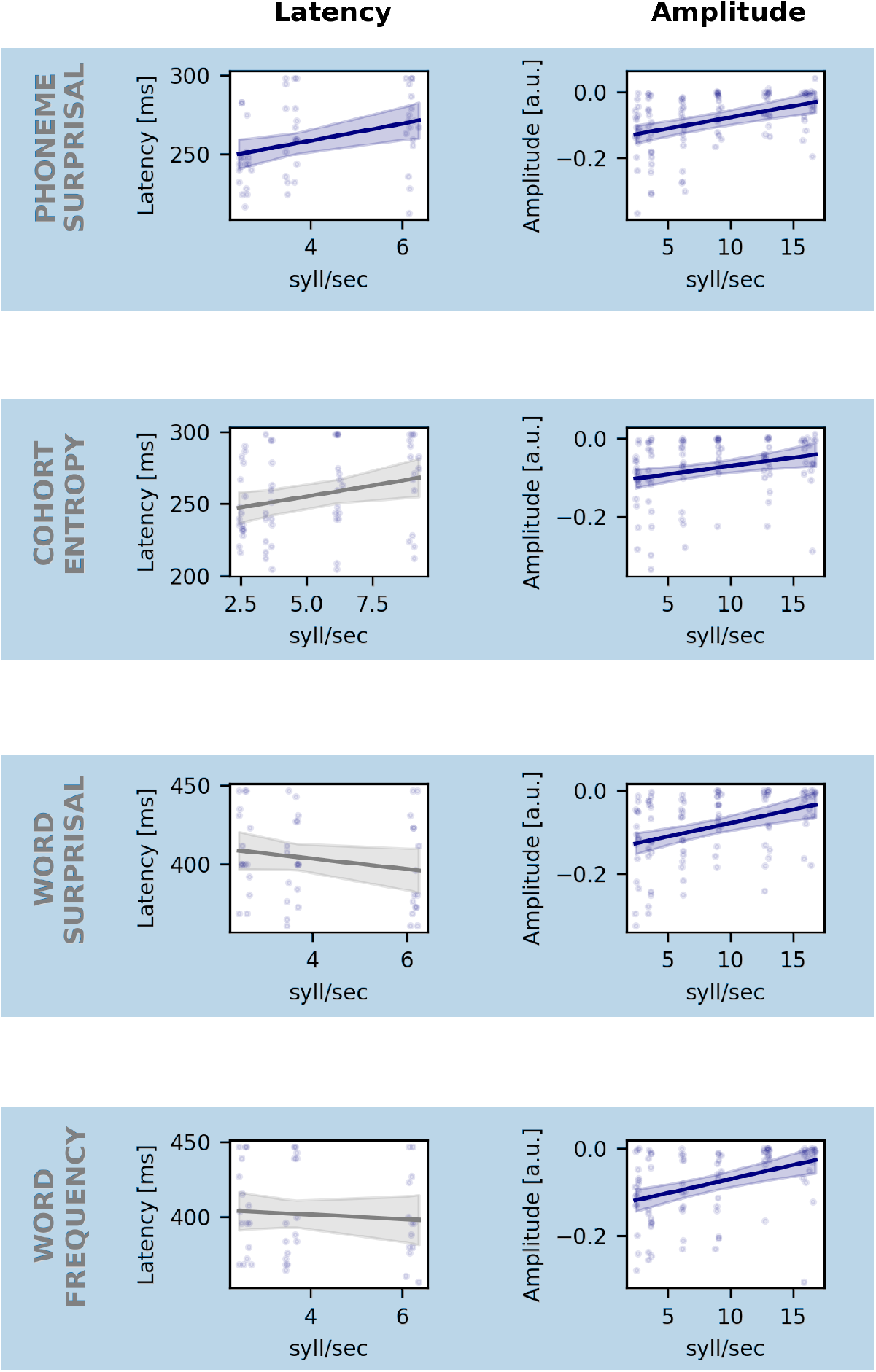
Effect of speech rate on the amplitude and latency of linguistic representation. Each row shows the effect of speech rate on latency (left plot) and the amplitude (right plot) of the neural response to phoneme surprisal (top row), cohort entropy (second row), word surprisal (third row) and word frequency (bottom row). The blue line shows the model’s prediction for each speech rate; the shaded area indicates the confidence interval of the model’s prediction. The non-significant models are shown in grey. Remark that we only include the latency of significant peaks for the latency analysis.

## 4 DISCUSSION

We aimed to investigate whether neural speech processing can capture the effect of gradually decreasing speech understanding by manipulating the speech rate. With increasing speech rate, we found that neural tracking of the acoustic features increased, while neural tracking of the linguistic features decreased.

### 4.1 Effect of speech rate on acoustic neural processing of speech

We hypothesized that neural tracking of acoustic features would increase with increasing speech rate because more acoustic information is present. We found an increase of acoustic neural tracking with increasing speech rate and thus decreasing speech understanding, confirming our hypothesis. When speech becomes faster, the model is better at predicting acoustic speech features.

This increase of acoustic neural tracking with increasing speech rate, and thus decreasing speech understanding, seems discrepant with previous research trying to link acoustic neural tracking to speech understanding using, for example, speech-in-noise paradigms (Vanthornhout et al., 2018; Verschueren et al., 2020; Ding and Simon, 2013; Iotzov and Parra, 2019; Etard and Reichenbach, 2019). The experimental paradigm could explain this discrepancy. Previous studies used, for example, noise to manipulate speech understanding. In those cases, decreased neural tracking was accompanied by a decrease in speech understanding and an acoustically degraded speech signal. Because speech understanding and signal-to-noise ratio are highly correlated, it is challenging to unravel to what extent the decreased neural tracking is driven by decreased speech understanding or the signal-to-noise ratio used to vary speech understanding. In this study, we manipulated speech understanding by speeding up the speech signal and preserving its signal-to-noise ratio, in contrast to the speech in noise studies. We hypothesize that the brain mainly responds to acoustic boundaries, i.e. onsets of sounds, which are more prominent in the faster speech presented in this study, explaining the increasing tendency. In addition, because the speech is sped up, the duration of the silences in between words or sentences inherently decreases, which increases the amount of speech data allowing the model to improve its estimate of the TRF and obtain higher prediction accuracies. However, this increase of acoustic neural tracking and speech rate is not linear: it saturates, and decreases again at very high speech rates. A first hypothesis is related to the motor cortex. When participants listen to the speech, they tend to mimic the speech in their brain, activating motor areas (Casas et al., 2021). However, most speakers cannot produce speech as fast as 16.2 syllables/sec. Hence, the corresponding mouth movements are unnatural, which implies that the listener cannot mimic the speech in their brain anymore, decreasing the related responses in the motor areas and thus brain responses to the acoustic speech features in general. A different hypothesis is related to the stimulus characteristics. When increasing the speech rate, the spectrogram and acoustic edges contain more and more peaks. Possibly, at very high speech rates, these peaks occur so fast after each other making it difficult for the brain to perceive them still (see Figure 1), similar to speech in noise studies where the amount of noise can mask acoustic boundaries. Therefore, it is difficult to attribute this decrease in acoustic neural tracking at very high speech rates (or low signal-to-noise ratios): a decrease in speech understanding or a decrease in neural detection of acoustic boundaries, or a combination of both?

To better understand the observed quadratic tendency of acoustic neural tracking with increasing speech rate, we investigated the TRFs of the speech features separately. Two significant peaks with opposite behavior could be observed for both acoustic features. The first peak is the largest. With increasing speech rate the amplitude of peak 1 increases, saturates and decreases again at very high speech rates. This is similar to the previously discussed neural acoustic tracking results. On the other hand, the second peak amplitude decreases with increasing speech rate. This discrepancy is intriguing as it suggests that both peaks have different underlying brain processes as confirmed by literature (Picton, 2011; Brodbeck and Simon, 2020). Peak 1 occurs relatively fast, around 50 ms, and is probably mainly related to the acoustics of the incoming speech and thus benefits from an increased speech rate. Peak 2, on the other hand, occurs somewhat later, around 150 ms, and could, in addition to the acoustics, be influenced by top-down processing related to speech understanding and attention (Ding and Simon, 2012; Vanthornhout et al., 2019). Besides the amplitude, we also investigated the latencies. The latency of the first peak increases with increasing speech rate. Increased latencies are often observed in more complex conditions with a higher task demand, like for example lower stimulus intensity, vocoded speech or speech in noise (Mirkovic et al., 2019; Verschueren et al., 2021; Kraus et al., 2020). The latency of the neural responses can also be related to neural processing efficiency (Bidelman et al., 2019; Gillis et al., 2021a). In more detail, a larger latency indicates that more processing time is required to process the same speech characteristics, showing reduced neural processing efficiency. More words and phonemes need to be processed as the speech rate increases, resulting in a more challenging condition to process the incoming speech.

### 4.2 Effect of speech rate on linguistic neural processing of speech

When speech becomes faster, speech understanding drops. Interestingly, this same decrease can be observed in linguistic neural tracking, in contrast to acoustic neural tracking (section 4.1). It is interesting to note that the pattern between speech rate and acoustic and linguistic neural tracking differs. Linguistic neural tracking seems linearly related to speech rate, while acoustic neural tracking shows non-linearity: it increases, stagnates and eventually slightly decreases for the highest speech rates. To the best of our knowledge, this is the first study that evaluates linguistic neural tracking when manipulating the level of speech understanding as a gradual effect. The studies of Brodbeck et al. (2018) and Broderick et al. (2018) using a two-talker paradigm are most comparable. They compared two conditions, i.e., intelligible and attended speech versus unintelligible and ignored speech, but not the spectrum in between. Nevertheless, their findings converge with our results and support our hypothesis of linguistic neural tracking as a neural marker of speech understanding. When the speech is not understood or ignored, the brain does not track the linguistic aspects of the speech, while for intelligible speech linguistic tracking is present.

To better understand the observed decrease of linguistic neural tracking with increasing speech rate, we investigated the TRFs of the speech representations separately. We observed a characteristic negative peak for each linguistic representation as observed in previous literature (e.g. Brodbeck et al., 2018; Gillis et al., 2021b; Weissbart et al., 2019). Although the responses are very small, they are significantly different from zero and the TRF patterns converge with the patterns observed in previous literature. For the phoneme-related features, phoneme surprisal and cohort entropy, this peak occurs around 250 ms. For the word-related features, word surprisal and word frequency, this peak occurs somewhat later, around 350 ms. The difference in timescale between both feature groups could be linked to the different speech processing stages (phonemes versus words) they represent (Van Canneyt et al., 2021). Regarding the topographies of these peaks, the understandable speech conditions are associated with a typical topography, similar to the classical N400 responses characterized by central negative channels. As speech becomes less understandable, i.e., the speech rate increases, the associated topography disappears.

For all speech features, the amplitude of this negative peak decreases with increasing speech rate until it disappears at speech rates as high as 9.0 tot 12.9 syllables/sec. Gillis et al. (2021b) already showed that these linguistic representations have an added value above and beyond acoustic and lexical representations. However, the authors did not compare intelligible to unintelligible speech. Here, we elegantly showed that as the speech becomes less understandable but remains audible and acoustically intact (in contrast to speech in noise studies or vocoder studies), the characteristic negative peak decreases and finally disappears. Altogether, our results suggest that these characteristic negative peaks to linguistic representations could be neural correlates of speech understanding.

### 4.3 Caveats of this study

As mentioned in the discussion, it is difficult to fully attribute a decrease in acoustic neural tracking at very high speech rates to a decrease in speech understanding (top-down) or a decrease in neural detection of acoustic boundaries (bottom-up). This is an important caveat in studies manipulating speech understanding by changing the acoustics of the speech signal. We cannot be sure whether the observed decrease is driven by speech rate or speech understanding. To enable a clean separation of acoustics and speech understanding, the speech acoustics should remain the same over conditions, such as in the paradigm used by Di Liberto et al. (2018). They use a priming paradigm where vocoded speech is presented before and after priming, i.e., learning the speech’s meaning by presenting the non-vocoded sentence before the vocoded sentence. This is an elegant way to disentangle acoustic from linguistic processing but only enables an on-off (understanding versus no understanding) analysis. In addition, priming is only feasible with a small number of words or sentences. Using a continuous story would make this task too complex for the participants.

The second caveat in this study is that, per condition, neural speech tracking is trained on the same amount of data in terms of time but not feature instances. For example, 10 minutes of slow speech contains fewer feature instances than 10 minutes of fast speech. In this study, we elected to keep the amount of EEG data constant because we know a model is affected by the amount of training data. Therefore, it is hard to conclude to what extent the increased acoustic neural tracking at higher speech rates is influenced by the increased number of feature instances in the model. Ideally, we would be able to compare both models. This is unfortunately not convenient in this study because 10 minutes of slow speech corresponds to only 94 seconds of speech in the fastest condition, which is not enough to train a model. However, as suggested by the reviewer, we did redo the analysis for the 4 lowest speech rates reducing the amount of EEG until each speech rate condition contained the same amount of feature instances. The effect of speech rate on acoustic tracking was similar for both analysis: when speech rate increases, acoustic neural tracking increases. The only difference is that this increase is faster when keeping the amount of data constant instead of preserving the amount of feature instances.

### 4.4 Conclusion

Using a speech rate paradigm, we map how the level of speech understanding affects acoustic and linguistic neural speech processing. When speech rate increases, acoustic neural tracking increases, although speech understanding drops. However, the amplitude of the later acoustic neural response decreases with increasing speech rate, suggesting influence of top-down processing related to speech understanding and attention. In contrast, linguistic neural tracking decreases with increasing speech rate and even disappears when speech is no longer understood. Altogether, this suggests that linguistic neural tracking could possibly be a more direct predictor of speech understanding compared to acoustic neural tracking.

## Acknowledgment

The authors would like to thank Sofie Keunen and Elise Verwaerde for their help in data acquisition.

## REFERENCES

Ahissar, E., Nagarajan, S., Ahissar, M., Protopapas, A., Mahncke, H., and Merzenich, M. M. (2001). Speech comprehension is correlated with temporal response patterns recorded from auditory cortex. Proceedings of the National Academy of Sciences of the United States of America 98, 13367–72. doi:10.1073/pnas.201400998

Baayen, R. H., Piepenbrock, R., and Gulikers, L. (1996). The celex lexical database (cd-rom)

Bates, D., Mächler, M., Bolker, B., and Walker, S. (2015). Fitting linear mixed-effects models using lme4. Journal of Statistical Software 67, 1–48. doi:10.18637/jss.v067.i01

Bidelman, G. M., Price, C. N., Shen, D., Arnott, S. R., and Alain, C. (2019). Afferent-efferent connectivity between auditory brainstem and cortex accounts for poorer speech-in-noise comprehension in older adults. Hearing research 382, 107795

Boursma, P. and Weenink, D. (2018). Praat: doing phonetics by computer [computer program]. version 6.0.37, retrieved 3 Februari 2018 from http://www.praat.org/

[Dataset] Brodbeck, C. (2020). Eelbrain 0.34. http://doi.org/10.5281/zenodo.3923991

Brodbeck, C., Hong, L. E., and Simon, J. Z. (2018). Rapid transformation from auditory to linguistic representations of continuous speech. Current Biology 28, 3976–3983

Brodbeck, C. and Simon, J. Z. (2020). Continuous speech processing. Current Opinion in Physiology 18, 25–31. doi:10.1016/j.cophys.2020.07.014

Broderick, M. P., Anderson, A. J., Liberto, G. M. D., Crosse, M. J., and Lalor, E. C. (2018). Electrophysiological Correlates of Semantic Dissimilarity Reflect the Comprehension of Natural, Report Electrophysiological Correlates of Semantic Dissimilarity Reflect the Comprehension of Natural, Narrative Speech. Current Biology 28, 803–809. doi:10.1016/j.cub.2018.01.080

Casas, A. S. H., Lajnef, T., Pascarella, A., Guiraud-vinatea, H., Laaksonen, H., Bayle, D., et al. (2021). Neural oscillations track natural but not artificial fast speech: Novel insights from speech-brain coupling using meg. NeuroImage 244, 118577

David, S. V., Mesgarani, N., and Shamma, S. A. (2007). Estimating sparse spectro-temporal receptive fields with natural stimuli. Network: Computation in Neural Systems 18, 191–212. doi:10.1080/09548980701609235

Decruy, L., Das, N., Verschueren, E., and Francart, T. (2018). The self-assessed Békesy procedure: validation of a method to measure intelligibility of connected discourse. Trends in Hearing 22, 1–13. doi:10.1177/2331216518802702

Di Liberto, G. M., Lalor, E. C., and Millman, R. E. (2018). Causal cortical dynamics of a predictive enhancement of speech intelligibility. NeuroImage 166, 247–258. doi:10.1016/j.neuroimage.2017.10.066

Ding, N. and Simon, J. Z. (2012). Emergence of neural encoding of auditory objects while listening to competing speakers. Proceedings of the National Academy of Sciences of the United States of America 109, 11854–9. doi:10.1073/pnas.1205381109

Ding, N. and Simon, J. Z. (2013). Adaptive Temporal Encoding Leads to a Background-Insensitive Cortical Representation of Speech. Journal of Neuroscience 33, 5728–5735. doi:10.1523/JNEUROSCI.5297-12.2013

Duchateau, J., Kong, Y. O., Cleuren, L., Latacz, L., Roelens, J., Samir, A., et al. (2009). Developing a reading tutor: Design and evaluation of dedicated speech recognition and synthesis modules. Speech Communication 51, 985–994

Elzhov, T. V., Mullen, K. M., Spiess, A.-N., and Bolker, B. (2016). minpack.lm: R Interface to the Levenberg-Marquardt Nonlinear Least-Squares Algorithm Found in MINPACK, Plus Support for Bounds. R package version 1.2-1

Etard, O. and Reichenbach, T. (2019). Neural speech tracking in the theta and in the delta frequency band differentially encode clarity and comprehension of speech in noise. Journal of Neuroscience 39, 5750–5759

Francart, T., van Wieringen, A., and Wouters, J. (2008). APEX 3: a multi-purpose test platform for auditory psychophysical experiments. Journal of Neuroscience Methods 172, 283–293

Gillis, M., Decruy, L., Vanthornhout, J., and Francart, T. (2021a). Hearing loss is associated with delayed neural responses to continuous speech. bioRxiv

Gillis, M., Vanthornhout, J., Simon, J. Z., Francart, T., and Brodbeck, C. (2021b). Neural markers of speech comprehension: measuring EEG tracking of linguistic speech representations, controlling the speech acoustics. Journal of neurocience, [Accepted for publication]

Gwilliams, L. and Davis, M. H. (2022). Extracting language content from speech sounds: the information theoretic approach. In Speech Perception (Springer). 113–139

[Dataset] Heeris, J. (2014). Gammatone filterbank toolkit 1.0. https://github.com/detly/gammatone

Hertrich, I., Dietrich, S., Trouvain, J., Moos, A., and Ackermann, H. (2012). Magnetic brain activity phase-locked to the envelope, the syllable onsets, and the fundamental frequency of a perceived speech signal. Psychophysiology 49, 322–334. doi:10.1111/j.1469-8986.2011.01314.x

Iotzov, I. and Parra, L. (2019). EEG can predict speech intelligibility To. J. Neural Eng. 16, 036008 (11p). doi:10.1088/1741-2552/ab07fe

Keuleers, E., Brysbaert, M., and New, B. (2010). Subtlex-nl: A new measure for dutch word frequency based on film subtitles. Behavior research methods 42, 643–650

Koskinen, M., Kurimo, M., Gross, J., Hyv, A., and Hari, R. (2020). NeuroImage Brain activity re fl ects the predictability of word sequences in listened continuous speech. NeuroImage 219, 116936. doi:10.1016/j.neuroimage.2020.116936

Kraus, F., Tune, S., Ruhe, A., Obleser, J., and Woestmann, M. (2020). Unilateral acoustic degradation delays attentional separation of competing speech. bioRxiv

Lesenfants, D., Vanthornhout, J., Verschueren, E., Decruy, L., and Francart, T. (2019). Predicting individual speech intelligibility from the neural tracking of acoustic- and phonetic-level speech representations. Hearing Research 380, 1–9. doi:10.1016/j.heares.2019.05.006

MacPherson, A. and Akeroyd, M. A. (2014). A method for measuring the intelligibility of uninterrupted, continuous speech. The Journal of the Acoustical Society of America 135, 1027–1030

Maris, E. and Oostenveld, R. (2007). Nonparametric statistical testing of eeg-and meg-data. Journal of neuroscience methods 164, 177–190

Mirkovic, B., Debener, S., Schmidt, J., Jaeger, M., and Neher, T. (2019). Effects of directional sound processing and listener’s motivation on eeg responses to continuous noisy speech: Do normal-hearing and aided hearing-impaired listeners differ? Hearing Research 377, 260–270

Müller, J. A., Wendt, D., Kollmeier, B., Debener, S., Brand, T., and Hunter, C. R. (2019). Effect of Speech Rate on Neural Tracking of Speech. Frontiers in psychology 10, 1–15. doi:10.3389/fpsyg.2019.00449

Nourski, K. V., Reale, R. A., Oya, H., Kawasaki, H., Kovach, C. K., Chen, H., et al. (2009). Temporal Envelope of Time-Compressed Speech Represented in the Human Auditory Cortex. The Journal of Neuroscience 29, 15564–15574. doi:10.1523/JNEUROSCI.3065-09.2009

Picton, T. W. (2011). Human Auditory Evoked Potentials (San Diego: Plural Publishing inc.)

Sanders, L. D. and Neville, H. J. (2003). An erp study of continuous speech processing: I. segmentation, semantics, and syntax in native speakers. Cognitive Brain Research 15, 228–240

Shannon, R. V., Zeng, F. G., Kamath, V., Wygonski, J., and Ekelid, M. (1995). Speech recognition with primarily temporal cues. Science 270, 303–304

Slaney, M. (1998). Auditory toolbox. Interval Research Corporation, Tech. Rep 10

Somers, B., Francart, T., and Bertrand, A. (2018). A generic EEG artifact removal algorithm based on the multi-channel Wiener filter. Journal of neural engineering 15. doi:10.1088/1741-2552/aaac92

Van Canneyt, J., Gillis, M., Vanthornhout, J., and Francart, T. (2021). Neural tracking as an objective measure of auditory perception and speech intelligibility. bioRxiv

Vanthornhout, J., Decruy, L., and Francart, T. (2019). Effect of task and attention on neural tracking of speech. BioRxiv, doi: http://dx.doi.org/10.1101/568204 doi:10.3389/fpsyg.2019.00449

Vanthornhout, J., Decruy, L., Wouters, J., Simon, J. Z., and Francart, T. (2018). Speech intelligibility predicted from neural entrainment of the speech envelope. JARO 19, 181–191. doi:10.1007/s10162-018-0654-z

Verschueren, E., Vanthornhout, J., and Francart, T. (2020). The effect of stimulus choice on an EEG-based objective measure of speech intelligibility. Ear & Hearing, Publish Ahead of preprintdoi:https://doi.org/10.1101/421727

Verschueren, E., Vanthornhout, J., and Francart, T. (2021). The effect of stimulus intensity on neural envelope tracking. Hearing Research 403, 108175

Weissbart, H., Kandylaki, K. D., and Reichenbach, T. (2019). Cortical Tracking of Surprisal during Continuous Speech Comprehension. Journal of cognitive neuroscience 32, 155–166. doi:10.1162/jocna01467

